# The effects of skin tone on photoacoustic imaging and oximetry

**DOI:** 10.1101/2023.08.17.553653

**Authors:** Thomas R. Else, Lina Hacker, Janek Gröhl, Ellie V. Bunce, Ran Tao, Sarah E. Bohndiek

## Abstract

**Significance:** Photoacoustic imaging (PAI) provides contrast based on the concentration of optical absorbers in tissue, enabling the assessment of functional physiological parameters such as blood oxygen saturation (sO_2_). Recent evidence suggests that variation in melanin levels in the epidermis leads to measurement biases in optical technologies, which could potentially limit the application of these biomarkers in diverse populations.

**Aim:** To examine the effects of skin melanin pigmentation on photoacoustic imaging and oximetry.

**Approach:** We evaluated the effects of skin tone in PAI using a computational skin model, two-layer melanin-containing tissue-mimicking phantoms, and mice of a consistent genetic background with varying pigmentations. The computational skin model was validated by simulating the diffuse reflectance spectrum using the adding-doubling method, allowing us to assign our simulation parameters to approximate Fitzpatrick skin types. Monte Carlo simulations and acoustic simulations were run to obtain idealised photoacoustic images of our skin model. Photoacoustic images of the phantoms and mice were acquired using a commercial instrument. Reconstructed images were processed with linear spectral unmixing to estimate blood oxygenation. Linear unmixing results were compared with a learned unmixing approach based on gradient-boosted regression.

**Results:** Our computational skin model was consistent with representative literature for *in vivo* skin reflectance measurements. We observed consistent spectral colouring effects across all model systems, with an overestimation of sO_2_ and more image artefacts observed with increasing melanin concentration. The learned unmixing approach reduced the measurement bias, but predictions made at lower blood sO_2_ still suffered from a skin tone-dependent effect.

**Conclusion:** PAI demonstrates measurement bias, including an overestimation of blood sO_2_, in higher Fitzpatrick skin types. Future research should aim to characterise this effect in humans to ensure equitable application of the technology.

## 1. INTRODUCTION

Photoacoustic imaging (PAI) is an emerging molecular imaging modality that provides contrast based on the absorption of light by different molecules (chromophores) in tissue. These molecules may be endogenous, such as haemoglobin, melanin, lipids and water, or exogenous, covering a range of contrast agents.^1, 2^ PAI has the potential to recover the concentration of each of these chromophores based on their unique optical absorption spectra, by acquiring data at multiple wavelengths and subjecting the reconstructed images to multispectral processing methods.^3^ For example, the recovery of relative concentrations of oxy- and deoxy-haemoglobin is commonly used to derive oxygenation maps from PAI with linear spectral unmixing.^4^ Obtaining accurate maps of chromophore concentrations, however, is limited by spatial variations in light fluence, which are significant at depth and are highly dependent on the surrounding distribution of optical absorbers and scatterers.^3^ This is often referred to as ‘spectral colouring’. Several factors can affect tissue optical properties, including age,^5^ low blood perfusion^6^ or skin tone,^7^ which consequently affect the ability to extract quantitative information from photoacoustic imaging.

Measurement biases introduced by the differential absorption of melanin in darker skin tones have been identified in several technologies where light passes through the skin before making a quantitative measurement, including pulse oximeters,^8–18^ bilirubinometers,^19, 20^ wearable technologies,^21–24^ cerebral oximeters^25^ and optical reflectance measurements.^26^ These biases can have significant implications for patient management, an issue that came to the fore in the management of COVID-19 patients, where the over-estimation of blood oxygenation by pulse oximetry in black patients may have led to under-diagnosis of hypoxaemia compared to white patients.^17^ Photoacoustic imaging also relies on the transmission of light through the skin and is thus also expected to be subject to similar limitations.

Skin tone is typically defined according to the Fitzpatrick scale, which provides a ranking based on the response of the skin to ultraviolet light determined by a questionnaire, from type I (always burns, never tans) to type VI (never burns, pigmented skin). A few studies have begun to emerge that examine skin tone measurement bias in PAI.^27–30^ Higher Fitzpatrick skin type grading was shown to reduce PAI signal-to-noise ratio deep in the tissue and introduced biases in the estimated blood oxygenation.^27–30^ An empirical calibration scheme has been proposed to correct these biases.^29^ These early studies highlight the key challenges associated with varying skin tones in PAI, how we measure skin tone, and how we develop fluence compensation schemes. The first of these key challenges could be tackled using more quantitative measures of skin tone, for example, measurements based on colourimeters. The second may be improved by data-driven approaches, which improve on standard linear unmixing schemes for photoacoustic oximetry, but have not been tested in cases of varying skin tone.^4, 31–35^

Here, we undertook a systematic evaluation of the effects of skin melanin concentration on PAI, using a combination of simulation, phantom, and animal models to examine the highlighted challenges. We illustrate that linear spectral unmixing is highly dependent on skin tone and introduces biases in photoacoustic oximetry. These biases mirror observations made in pulse oximetry.^17^ We demonstrate that learned spectral decolouring can partially compensate for this bias, recovering less biased oxygenation values. We underscore these findings in animal studies, showing a systematic overestimation of blood oxygenation with increasing skin pigmentation. Our results provide a theoretical basis upon which we can interpret future studies in patients with different skin tones.

## 2. METHODS

### 2.1 Computational models

To assess the effects of skin tone on photoacoustic imaging in a controlled setting, we developed a computational forearm model imitating the structure and physical properties of the skin. The model was constructed in Python using absorption and reduced scattering spectra obtained from the literature.^36^ Three variations of the model were used: a layer-only model for calibration with adding-doubling reflectance simulations, a varying blood oxygenation model for optical modelling, and a realistic model for full photoacoustic simulation. These distinctions were made to allow thorough characterisation of the models under computational constraints. To bridge the gap between simulation and experiment, we also simulated optical absorption in the agarose phantom model described in Section 2.5.

The first model consisted of three parallel layers. The first layer represented the epidermis, modelled as 60 µm thick, with optical absorption from a varying concentration of melanosomes. The melanosome volume fraction was chosen by first simulating diffuse reflectance (Section 2.2) over a range previously identified in the literature (1.3 % to 43 %^37^), and adjusting the simulation range to align as closely as possible with individual typology angle (ITA) values associated with different Fitzpatrick types.^38^ The resulting parameters were then qualitatively verified using a publicly available integrating sphere forearm skin reflectance dataset (Figure S1).^39^ We settled on six logarithmically-spaced values of melanosome concentration between 2 % v/v and 40 % v/v, representing the six Fitzpatrick skin types. The second layer, representing the dermis, was 1 mm thick, containing blood (0.2% v/v, 70% oxygenation), water (65% v/v) and lipid (20% v/v).^36^ The third layer, representing generic tissue, filled the rest of the simulation volume. It consisted of blood (2.5% v/v, 70% oxygenation), water (65% v/v) and lipid (20% v/v). The absorption spectra for these three layers are shown in Figure 1 A. Scattering properties were defined by a combination of Rayleigh and Mie scattering, assuming a constant scattering anisotropy of 0.9. These values are representative of typical values in each region.^36^ This simplified layer model was used for validation purposes, using adding-doubling simulations of diffuse reflectance. The structure formed the basis of all simulations, with blood vessels added for Monte-Carlo simulations.

**Figure 1.**
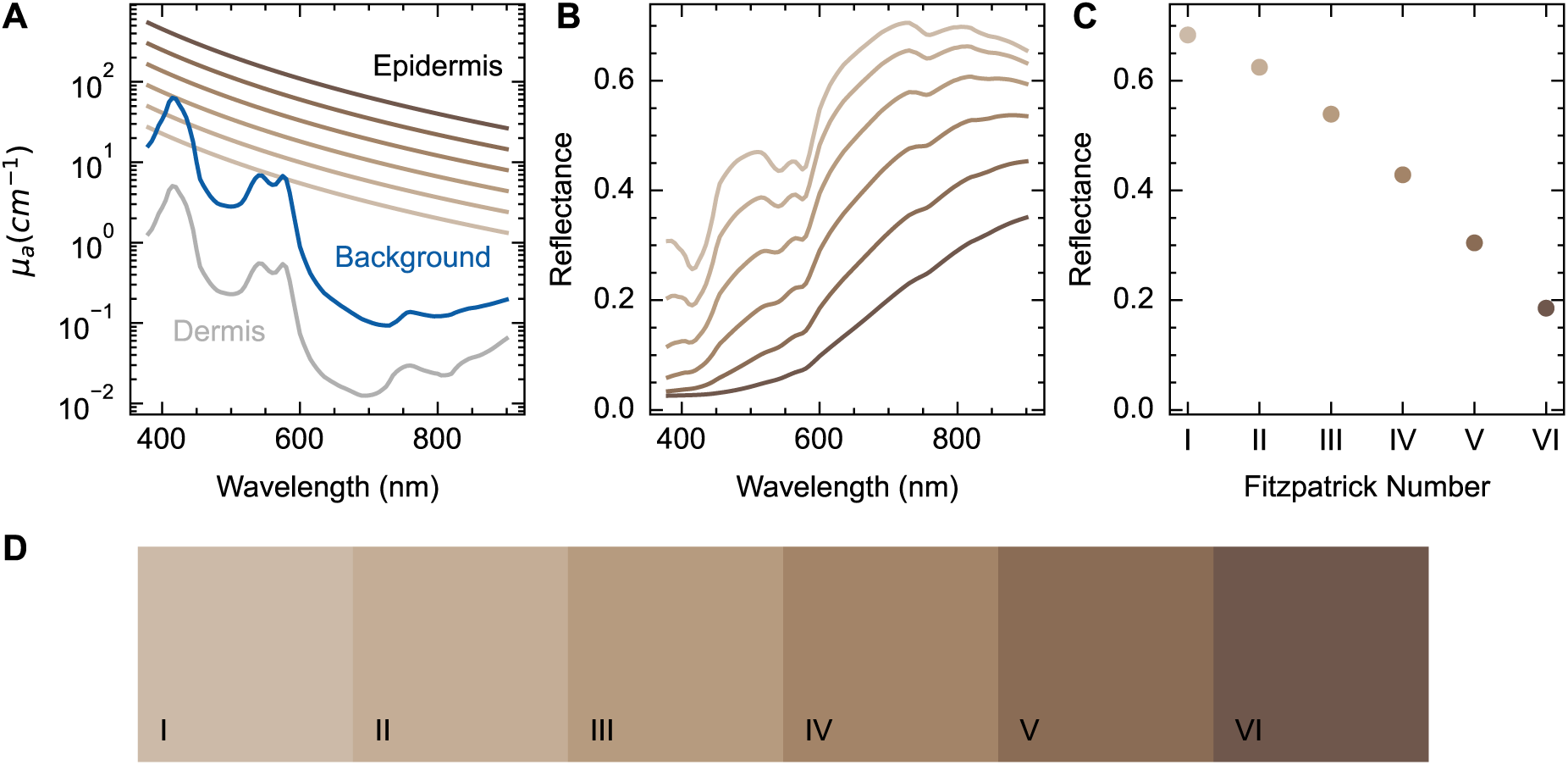
Diffuse reflectance simulations validate Monte-Carlo simulation parameters and enable the assignment of simulations to equivalent Fitzpatrick skin types. (A) Optical absorption spectra in the tissue model were assigned according to the desired melanosome concentration (2 % to 40 % v/v, gradations of brown) in the skin layer, dermis below (grey) and background absorber (blue). (B) Diffuse reflectance spectra were calculated using the adding-doubling method. (C) Reflectance values simulated at 685 nm cover the physiological range of skin reflectance. (D) Red, green and blue images were simulated from the reflectance spectra and the six models were assigned to the six Fitzpatrick skin types (shown by inset label I - VI).

To model the effects of changing blood oxygenation in a blood vessel on the photoacoustic initial pressure, Monte-Carlo simulations were run with the varying blood oxygenation model, giving the optical absorption distribution. This model was based on the layer structure described above with the addition of a cylinder of radius 1.25 mm, at a depth of 2.5 mm below the epidermis, representing a blood vessel of varying oxygenation (between 0 % and 100 %).

To evaluate the effects of a more realistic tissue model, a third model, the realistic forearm model, was subjected to both optical and acoustic simulations, giving full photoacoustic data that was processed using normal reconstruction and analysis procedures. Two cylinders were added to the layer model, one representing an artery (100 %-oxygenated blood, radius 1.25 mm at a depth of 2.5 mm below the epidermis) and one representing a vein (70 % oxygenated blood, radius 0.625 mm at a depth of 2.5 mm).

We also ran optical simulations of the cylindrical phantoms (Section 2.5), with approximately identical optical absorption and scattering properties as the agarose phantoms. We defined the model in Python, with a grid size of 100 µm in an array of size (220 *×* 220 *×* 220). The tube in the centre of the phantom was modelled as a cylindrical absorber of radius 0.75 mm containing blood. Concentric absorbers were added representing the inner part of the phantom (23 mm diameter, water absorption and intralipid scattering) and the outer layer of the phantom (1 mm thick, varying melanosome volume fraction, intralipid scatterer). The absorption coefficients of the melanin layer were defined by finding the melanosome volume fraction that gave the same optical absorption coefficient at 700 nm as measured in the double integrating sphere system (Supplementary Table S1, Section 2.6).

### 2.2 Adding-doubling simulations

The layer-only skin model was validated using the adding-doubling model of diffuse reflectance^40^ to allow for fast evaluation of the reflectance spectra at several wavelengths (450 nm to 900 nm in 20 nm steps) in the visible and near-infrared ranges. We additionally calculated the reflectance at 685 nm to enable direct comparison with literature values.^41^ The reflectance spectra were converted into an RGB image and an individual typology angle (ITA) metric using a D65 standard light source and a 10-degree standard observer in the Python colour-science library.^42^ ITA has been proposed as a quantitative measure of skin tone and can be obtained from low-cost colourimeters. ITA was calculated from skin reflectance spectra using the following equation:

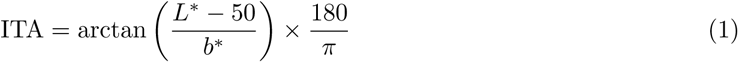

where *L^∗^* and *b^∗^* are taken from the CIELAB colour space, representing luminance (ranging from black (0) to white (100)) and the range from yellow to blue respectively.

### 2.3 Monte-Carlo simulations

We simulated the PAI optical forward model using the varying blood oxygenation model described above using a custom version of MCX (https://github.com/IMSY-DKFZ/mcx), giving initial pressure distributions that correspond to an idealised photoacoustic image. The model was defined on a grid with a pixel size of 60 µm, which allowed the simulation to run in a sufficiently short time while ensuring that the simulations were sufficiently detailed. The light transport through the model was simulated with 10^7^ photons per simulation, at 21 equally-spaced wavelengths between 700 nm and 900 nm inclusive.^43^

We used an illumination geometry which approximates a clinical PAI system (MSOT Acuity, iThera Medical GmbH, Munich, Germany), consisting of a single optical fibre with a beam divergence of 8.66*^°^*, 43.2 mm above the surface of the probe, directed at an angle of 22.4*^°^* from the imaging plane, intersecting the imaging plane 2.8 mm outside of the probe.^44^ Optical simulations of the agarose phantoms were run using the same customised version of MCX with an illumination geometry approximating a preclinical photoacoustic system (iThera MSOT inVision 256-TF, iThera Medical GmbH, Munich Germany), defined on a grid of size 100 µm. The illumination geometry consisted of five radially-distributed fibre bundle pairs.^44^ The version of MCX used in these experiments is available on GitHub: https://github.com/BohndiekLab/melanin-phantom-simulation-paper. 10^7^ photons were simulated per fibre bundle, for a total of 10^8^ photons. All simulations were run on a computer with 2 NVIDIA Quadro 8000 GPUs (48 GB GPU RAM each), Intel Xeon Gold 6230 CPU at 2.10 GHz (40 cores), and 256 GB RAM.

### 2.4 Acoustic simulations

To determine the impact of acoustic modelling, we ran the forward acoustic model in j-Wave^45^ to calculate the pressure time series detected by the ultrasound transducer array for the initial pressure distributions. Simulations were run in three dimensions, with a grid size of 240 µm, and a speed of sound of 1500 m s*^−^*^1^. Three-dimensional initial pressure distributions were taken from Monte-Carlo simulations and down-sampled by a factor of four, and their propagation was simulated. The resulting pressure distributions were evaluated using band-limited interpolation sensors in the position of the sensors in the photoacoustic system. The pressure time series were recorded and processed in the same way as the experimental data.

### 2.5 Phantom design

Phantoms were used to validate the simulations in a real photoacoustic system. A custom-made 3D-printed polylactic acid (PLA) mould was constructed to fabricate a two-layer cylindrical phantom with an inner scattering compartment and an outer compartment containing melanin to model the epidermis. The mould consisted of two cylindrical compartments, one of radius 9 mm and one of radius 10 mm. A polyvinyl chloride (PVC) tube (inner diameter 1.5 mm, outer diameter 2.1 mm; VWR 228-3857) was placed in the centre of the phantom mould to enable blood flow through the phantom.

Agarose-based phantoms were fabricated using standard methods.^46^ The inner phantom mixture consisted of a mixture of agarose for mechanical robustness and intralipid for scattering. 1.5 % (w/v) agarose (Sigma-Aldrich 05039) was added to Milli-Q water and heated until it dissolved. Once the mixture had cooled to approximately 40 °C, 2.08 % (v/v) of pre-warmed intralipid (20 % emulsion, Sigma-Aldrich I141) was added and mixed, before being poured into the inner compartment of the phantom mould and allowed to set, constituting the inner 9 mm radius of the phantom. Synthetic melanin (Sigma, M8631, 2.5 mg mL*^−^*^1^ stock in dimethyl sulfoxide) was added to a separate batch of the base mixture at the desired concentrations (Supplementary Table S1). Once the inner compartment was set, the outer layer mixture was added and again allowed to set at room temperature (Supplementary Figure S2).

### 2.6 Phantom characterisation

A custom double integrating sphere (DIS) system was used to characterise the optical absorption and reduced scattering coefficients of the phantom material. Samples were placed between two integrating spheres (Avantes, AvaSphere-50, 50 mm internal diameter), both of which were connected to a spectrometer (Avantes, Starline Avaspec-2048) via an optical fibre. The reflectance integrating sphere was connected to a broadband light source (Avantes, Avalight-HAL-s-mini) via an optical fibre. Circular samples (approximate diameter: 5 cm) of the agarose mixture with and without melanin were prepared with a thickness of approximately 3 mm. Sample thickness was measured using vernier callipers. Each agarose sample was placed between the two spheres and the transmittance and reflectance were measured. Optical scattering and absorption coefficients were calculated using the inverse adding doubling (IAD) method,^47^ assuming a scattering anisotropy factor of 0.89 and refractive index of 1.34.^48^ These measurements were made on three replicates of each sample, over a wavelength range of 450 nm to 900 nm at 1 nm steps.

### 2.7 Blood-flow circuit

A blood-flow circuit was used to control the oxygenation of blood flowing through the tube centred in the two-layer cylindrical phantom, enabling photoacoustic measurements of blood with known ground-truth blood oxygenation as described previously.^49, 50^ Briefly, we measured the oxygen partial pressure (pO_2_) and the photoacoustic spectrum of the blood simultaneously. We used fresh whole blood from Wistar rats (6 to 9 months, Charles River Laboratories). All blood samples were anti-coagulated in ethylenediaminetetraacetic acid (EDTA), stored at 4 *^°^*C, and processed within 48 h. The blood was pumped around the circuit using a peristaltic pump (Fisher Scientific CTP100), and the oxygenation was controlled by initially oxygenating the blood using hydrogen peroxide (30 % (w/w) in deionized water, Sigma-Aldrich 7722-84-1) and then slowly injecting 0.03 % w/v sodium dithionite (ACROS Organics 7775-14-6, in PBS) using a syringe pump (10 µL min*^−^*^1^, Harvard, MKCB2159V). To measure the blood pO_2_, two oxygen fluorescence quenching needle probes (Oxford Optronix, NX-BF/O/E) were placed before and after the phantom; the average of these measurements was taken as the ground-truth pO_2_ in the phantom. pO_2_ was converted to blood oxygenation (sO_2_) using the Severinghaus equation.^51^ The values were extracted using custom code on an Arduino UNO and fed to a computer over USB where they were read and stored using custom code in MATLAB. Experiments were conducted at 37 *^°^*C.

### 2.8 Animal procedures

All animal procedures were conducted under personal and project licences issued under the United Kingdom Animals (Scientific Procedures) Act, 1986 and reviewed by the Animal Welfare and Ethical Review Board at the CRUK Cambridge Institute. The protocols were approved locally under compliance forms CFSB2317 and CFSB2348.

To understand the effects of skin pigmentation in a model *in vivo* setting, we compared non-pigmented albino mice (n = 9, C57BL/6 albino, Janvier Labs, France, 10 to 13 weeks old) to pigmented “B6” mice (n = 10, C57BL/6J, Charles River, 10 to 13 weeks old). The B6 mice were further subdivided qualitatively based on the visual appearance of pigmentation patches in colour photographs (Supplementary Figure S3) into a non-pigmented group (n = 8 scans) and a pigmented group (n = 11 scans). Albino mice were imaged once and B6 mice were imaged twice (2 weeks apart) to allow the development of pigmentation after hair removal. Repeat measurements of the same mouse were treated as independent measurements for statistical analysis given the time lapse between measurements.

### 2.9 Photoacoustic imaging

Phantoms and mice were imaged using a pre-clinical multispectral photoacoustic tomography system (MSOT inVision-256TF, iThera Medical GmbH).^46^ Briefly, excitation pulses were provided by a tunable optical parametric oscillator, pumped by a nanosecond Nd:YAG laser (10 Hz repetition rate up to 7 ns pulse duration). A custom optical fibre assembly creates a diffuse ring of uniform illumination over the imaging plane within the sample. The sample is coupled to the transducers using a water bath. For ultrasound detection, an array of 256 toroidally focussed ultrasound transducers covering a 270*^°^* circular arc and radius of 4 cm was used with a centre frequency of 5 MHz and 60 % bandwidth.

For phantom imaging, a single cross-sectional slice of the phantom was imaged continuously during the blood deoxygenation procedure until the blood was completely deoxygenated. Phantoms were placed in a custom animal holder (iThera Medical GmbH, Munich, Germany), wrapped in a thin polyethylene membrane, with water used to couple the phantom to the membrane. Imaging was performed at wavelengths from 700 nm to 900 nm inclusive with 10 nm steps and frame averaging was enabled (10 averages) to improve the signal-to-noise ratio.

Mice were prepared for imaging according to our standard operating procedure.^46^ Briefly, the mice were shaved and depilatory cream was used to remove excess hair. The mice were anaesthetised using *<*3 % isoflurane in a 50 % pure oxygen/medical air mixture and placed in a custom animal holder (iThera Medical GmbH, Munich, Germany), wrapped in a thin polyethylene membrane, with ultrasound gel (Aquasonic Clear, Parker Labs) used to couple the skin to the membrane. The holder was then placed within the PAI system and immersed in deionised water maintained at 36 *^°^*C for Hb and HbO_2_ imaging acquisition. The animal respiratory rate was maintained in the range 70–80 b.p.m. with 1.5 % isoflurane concentration for the entire scan. A single cross-sectional slice through the kidneys and spleen was imaged. Frame averaging was enabled (4 averages) to minimise the effects of breathing motion artefacts. 10 repeat scans were taken at each of the following wavelengths 700 nm, 730 nm, 750 nm, 760 nm, 770 nm, 800 nm, 820 nm, 840 nm, 850 nm and 880 nm.

### 2.10 Image reconstruction and processing

Simulated, phantom and *in vivo* photoacoustic data were processed using the Python photoacoustic tomography analysis toolkit (PATATO, https://github.com/bohndieklab/patato).

Acoustic simulation images were reconstructed by filtered backprojection with a grid spacing of 60 µm and a field of view of 18 mm *×* 40 mm. Phantom and *in vivo* images were reconstructed by filtered backprojection with a grid spacing of 75 µm and a field of view of 25 mm. A band-pass filter was applied to all of the time series data, with a bandpass range of 5 kHz to 7 MHz.

Linear spectral unmixing was applied to all simulated, phantom and *in vivo* data sets using the matrix pseudo-inverse function in NumPy^52^ and literature values for the optical absorption spectra of oxy- and deoxy-haemoglobin.^36^ Photoacoustic estimates of the blood oxygenation (denoted with a superscript EST sO^EST^ to differentiate it from the true blood oxygenation, sO_2_) were made pixel-wise from the unmixing components as the ratio of the oxyhaemoglobin to the total haemoglobin. To evaluate the potential for a machine-learning approach to improve photoacoustic oximetry in the presence of melanin, a learned spectral decolouring^31^ algorithm was trained on a numerical tissue model containing varying levels of melanin in the epidermis layer (0.1% - 5% melanosome v/v). We simulated 500 unique volumes 19.2 mm^3^ in volume with a resolution of 0.3 mm that contained up to nine blood vessels with a random radius and blood oxygenation. The tissue background contained a blood concentration of 1 % (v/v) and an oxygenation of 70 %. These design parameters were developed independently of this study and simulations were obtained using the SIMPA framework.^44^ From the simulated initial pressure distributions, we used a histogram-based gradient boosting regression tree as the inversion algorithm with a maximum depth of 16 and otherwise default hyperparameters as implemented in the scikit-learn (version 1.2.1) Python package.

Unless otherwise specified, manual polygon regions of interest (ROIs) were drawn and average spectra or unmixed components were calculated by masking the defined regions. In phantoms, the ROIs were taken around the blood tube and the melanin compartment. The melanin compartment was defined by drawing an ROI around the whole phantom and eroding the polygon by 1 mm. The difference mask between the drawn and eroded polygons was applied to the photoacoustic images, giving the multispectral data within the melanin region. Mean photoacoustic and unmixed component values were extracted. In mice, ROIs were drawn around the whole mouse body. The body ROI was eroded by 1 mm to subdivide it into a “skin” region (outer 1 mm) and a “body” region (inner region). Negative pixel values were included in the median spectra calculation to illustrate image reconstruction artefacts. For unmixed sO^EST^ calculation, pixels were excluded if either of the unmixed oxy- or deoxy-haemoglobin coefficients were negative. Pixels near the open region of the detector were manually excluded from the analysis as the image reconstruction is less well-posed in that region, leading to artefacts.

Line plots of the photoacoustic signal through the mice were generated by constructing a line through the kidneys and vena cava of the mice. To ensure constant placement of this line, it was constructed to be 6.5 mm from the surface of the mouse closest to the spline, and perpendicular to an axis through the spine and aorta. This construction was observed to pass through the kidneys and vena cava in albino and non-pigmented mice. The same construction was applied to the heavily pigmented mice, where the kidneys and spleen were not visible, to give a direct comparison. The values of the reconstructed image, total haemoglobin map and blood oxygenation were calculated along this line using the interpnd function of SciPy.^52^

### 2.11 Statistics

Statistical analyses were conducted in Python using the statsmodels library. The relationship between continuous variables was quantified by linear regression. A log transformation was applied where specified to ensure that the residual normality assumption is upheld in each linear model. The residual normality assumption was tested using the Shapiro-Wilk test (p *>* 0.1 for all models). For the *in vivo* data, the relationship between each response variable and the mouse pigmentation was assessed using a linear model, with both mouse strain (Albino or B6) and pigmentation (not pigmented or pigmented) as independent variables. The model related the response variable (sO_2_ or photoacoustic signal) to a sum of the effect due to mouse strain and pigmentation with no interaction term.

## 3. RESULTS

### 3.1 The computational model of skin shows good agreement with literature based on adding-doubling reflectance measurements

We first validated our computational model of human skin, with the assigned optical absorption spectra in the epidermis (Figure 1 A), by calculating diffuse reflectance spectra using the adding-doubling method. We chose six values of melanosome concentration to represent the typical physiological range observed in the population. The calculated spectra show decreasing reflectance with increasing melanosome concentration and broadly increasing reflectance with wavelength, as one would expect given the profile of optical absorption of melanin (Figure 1 A, B). We then calculated the individual typology angle (ITA) for each reflectance spectrum, allowing direct comparison with *in vivo* measurements from skin colourimeters. These ITA values (67*^°^* to *−*47*^°^*) are consistent with previous observations made *in vivo* using the Fitzpatrick skin type (Table 1).^53^ Reflectance at 685 nm was evaluated for each tissue model, giving a range of reflection coefficients from 0.20 to 0.72 over our simulated range of melanosome concentrations (Figure 1 C, Table 1) again consistent with previous literature.^41^ To enable qualitative evaluation of our skin model, the reflection coefficients were converted into red, green and blue (RGB) images (Figure 1 D, Table 1). Our results align closely with literature values made of skin reflectance using a single integrating sphere (Supplementary Figure S1).^39^ For the remainder of the paper, results from different Fitzpatrick skin types are colour coded according to the calculated RGB values for consistency (Figure 1 D).

**Table 1.** Parameters of the skin simulations (melanosome concentration, absorption coefficient) and quantities derived from adding-doubling simulations (reflectance, individual typology angle, HEX colour value). The equivalent Fitzpatrick skin type was assigned based on literature values.^53^

| Melanosome concentration (v/v, %) | Absorption coefficient at 685 nm ($\text{cm}^{-1}$) | Reflectance at 685 nm | Individual typology angle ($^{\circ}$) | Assigned Fitzpatrick skin type | HEX colour |
| --- | --- | --- | --- | --- | --- |
| 2.0 | 3.45 | 0.72 | 67 | I | #CBB9A8 |
| 3.6 | 6.28 | 0.65 | 56 | II | #C3AC95 |
| 6.6 | 11.43 | 0.57 | 41 | III | #B69A7F |
| 12.1 | 20.81 | 0.46 | 20 | IV | #A28367 |
| 22.0 | 37.89 | 0.33 | -6.9 | V | #8A6B54 |
| 40.0 | 68.98 | 0.20 | -47 | VI | #6E574C |

### 3.2 Photoacoustic simulations with the skin model show spectral colouring due to skin pigmentation

Multispectral Monte-Carlo simulations using the computational skin model with an embedded blood vessel showed how the optical absorption values are affected by melanosome concentration and hence Fitzpatrick skin type (Figure 2 A). In simulations of fully oxygenated blood, we observed an increase in the optical absorption within the skin at all wavelengths with increasing Fitzpatrick type (Figure 2 B). The optical absorption in the epidermis increased more than 10-fold between Fitzpatrick I and Fitzpatrick VI at 700 nm (379 a.u. in FP I, 4128 a.u. in FP VI, Figure 2 C). Spectral colouring is observed within the blood vessel region, which manifests as a decrease in the mean-normalised absorption at lower wavelengths and a corresponding increase at higher wavelengths (Figure 2 D). Absorption measurements within the blood vessel region, however, decrease by a factor of 1.7 from Fitzpatrick I to Fitzpatrick VI (44.6 a.u. in FP I, 26.7 a.u. in FP VI, Figure 2 E). Combined optical and acoustic simulations of a realistic tissue model revealed additional image reconstruction artefacts associated with backprojection, including negative pixels and streaking artefacts (Supplementary Figure S4). Quantitative results from the blood and skin regions in the acoustic simulations were consistent with those made with solely optical simulations, so the remainder of the simulation study was performed using the Monte-Carlo optical simulation alone for computational practicality.

**Figure 2.**
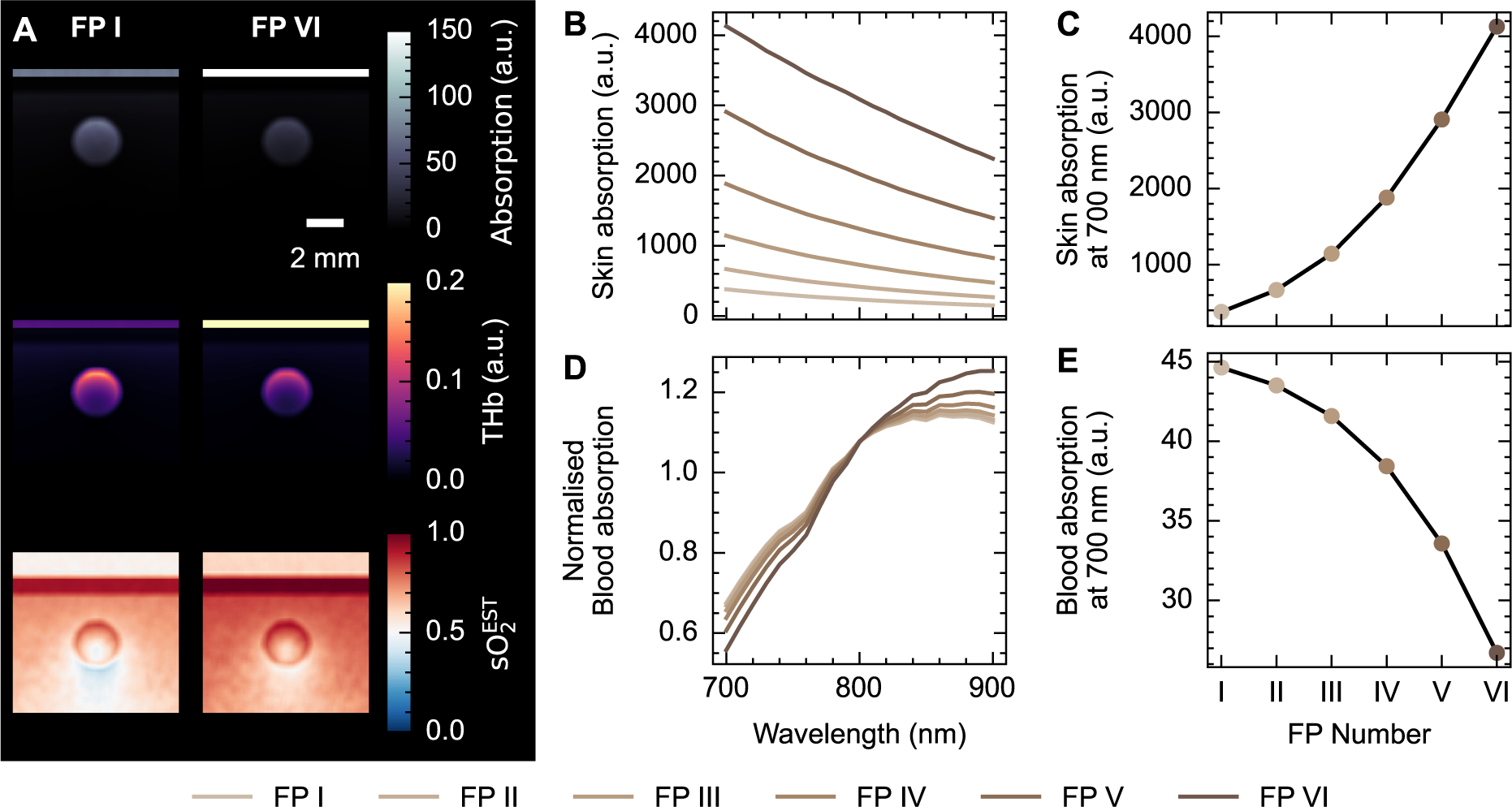
Monte-Carlo simulations of light transport through the skin model show decreased signal and increased spectral colouring in an underlying blood vessel with increasing Fitzpatrick skin type. (A) Photoacoustic initial pressure (upper), unmixed total haemoglobin (THb, middle), and unmixed blood oxygenation (sO^EST^, lower) in representative examples for fully oxygenated blood in the vessel. (B) Epidermis PA initial pressure as a function of wavelength for all Fitzpatrick types and (C) at 700 nm as a function of Fitzpatrick type. (D) Normalised blood PA initial pressure as a function of wavelength for all Fitzpatrick types and (E) at 700 nm as a function of Fitzpatrick (FP) type. Fitzpatrick types are denoted with graded brown colouration according to the RGB definitions in Figure 2 D. Scale bar = 2 mm applies to all images in (A).

Spectral colouring observed in the blood vessel biased the linear spectral unmixing blood oxygenation estimate (Figure 3 A). Encouragingly, the bias can be somewhat overcome when using learned spectral unmixing trained on independently generated numerical tissue models containing varying melanin concentrations (Figure 3 B). Considering the spectral changes observed (Figure 2 D, E), it appears that learned unmixing can better cope with the spectral corruption that presents as a reduction in normalised optical absorption at low wavelengths and a relative increase at higher wavelengths for higher Fitzpatrick skin types. With fully oxygenated blood, linear unmixing estimates of photoacoustic sO^EST^ reveal a non-linear relationship with Fitzpatrick skin type (Figure 3 B) and a linear relationship with melanosome concentration (p *<* 0.001, gradient = 0.218, 95% CI [0.200, 0.235]). Blood oxygenation estimated with learned unmixing also increases linearly with melanosome concentration, but with a smaller gradient (p *<* 0.001, gradient = 0.150, 95% CI [0.117, 0.183]) (Figure 3 D).

**Figure 3.**
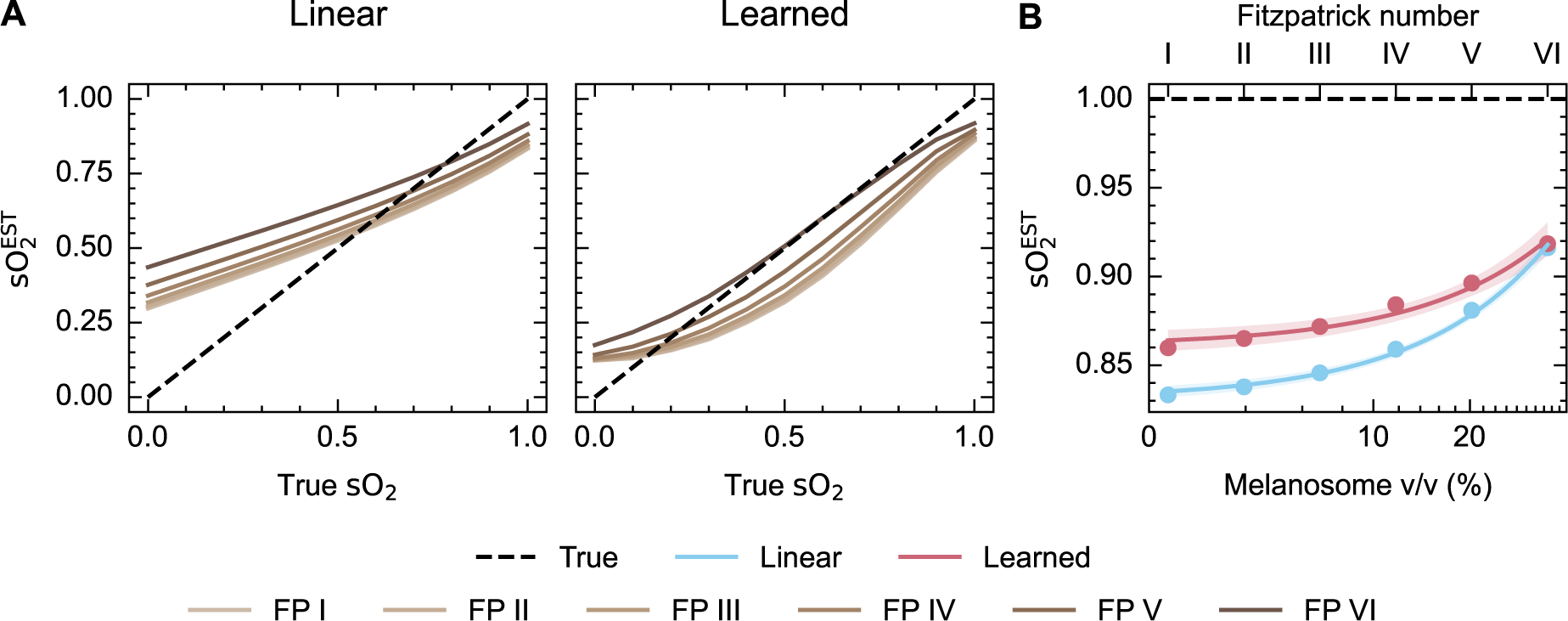
Estimation of blood oxygenation in simulated data using linear and learned spectral unmixing. (A) sO^EST^ estimates increase with increasing Fitzpatrick type when using linear spectral unmixing, while estimates are closer to the ground truth when using learned unmixing, particularly at high and low blood oxygenation levels. (B) sO^EST^ estimates for fully oxygenated blood using both approaches are shown as a function of Fitzpatrick type and melanosome volume fraction. The shaded area shows 95 % confidence intervals.

### 3.3 Blood flow phantoms confirm melanin-associated bias in an experimental photoacoustic system

Cylindrical phantoms were enclosed in a melanin-containing outer layer with concentrations equivalent to Fitzpatrick types below I, II and IV (Figure 4 A). A blood flow circuit was connected to a tube through the centre of the phantom, allowing continuous monitoring of the blood oxygenation. The optical properties of the phantoms were confirmed using double integrating sphere measurements (Supplementary Figure S5, Supplementary Table S1). By imaging the phantoms over time, as the blood was adjusted from fully oxygenated to fully deoxygenated, the photoacoustic measurements were related to the known blood oxygenation (Supplementary Figure S6). Images of these phantoms show strong absorption in the melanin outer layer and blood flow tube, with image reconstruction artefacts visible around the blood inclusion and near the melanin layer, particularly visible with the higher concentrations of melanin (Figure 4 B). The photoacoustic signal in the outer layer increases with melanin concentration (Figure 4 B, C). Wavelength-dependent spectral colouring is observed in the blood spectrum with increasing melanin concentration, with normalised photoacoustic signals relatively lower at shorter wavelengths and higher at longer wavelengths, consistent with the optical absorption properties of melanin (Figure 4 D) and simulation findings.

**Figure 4.**
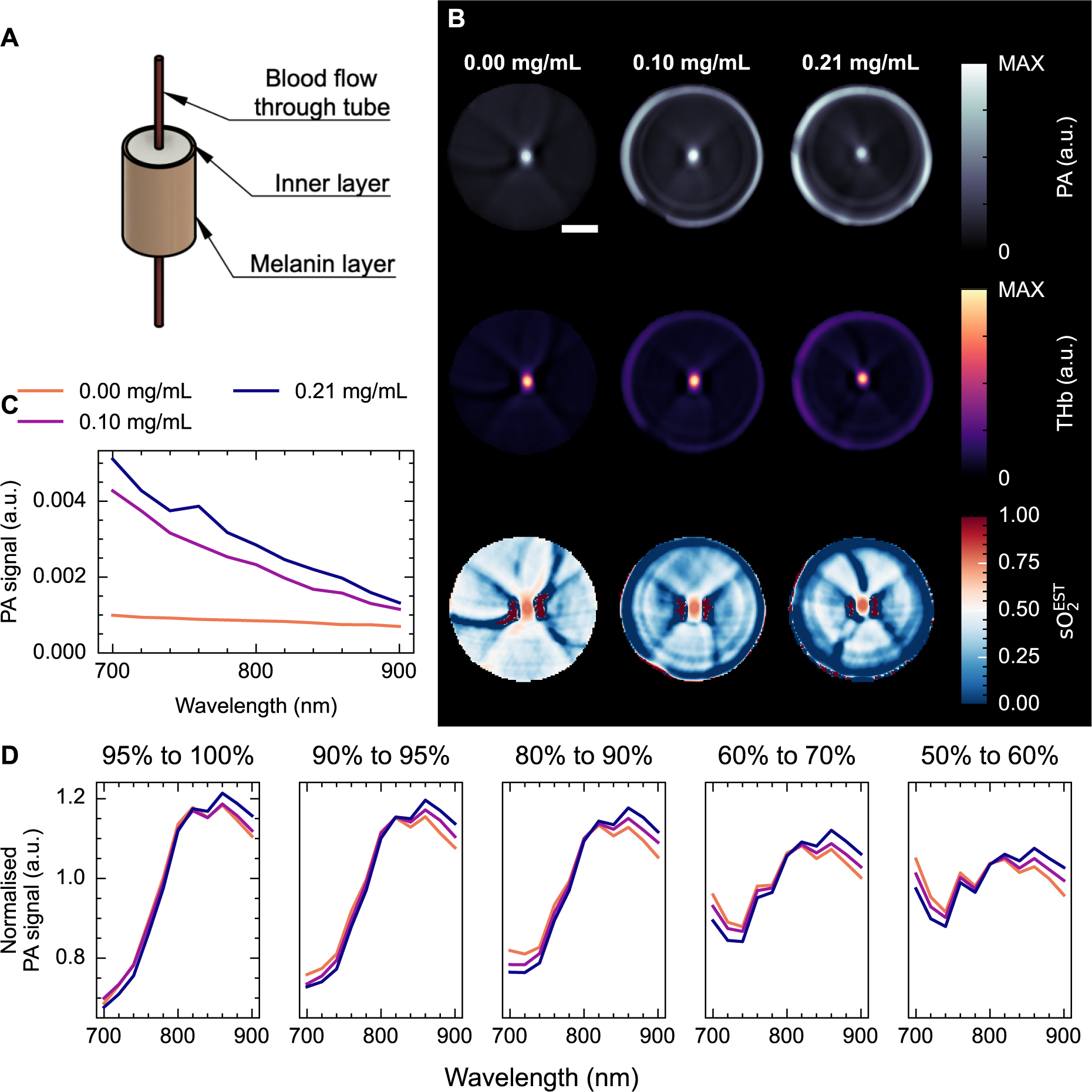
Skin-mimicking blood-flow phantoms show spectral colouring. (A) Schematic diagram of the agarose phantom setup, showing a 1 mm-thick outer layer containing synthetic melanin and a blood inclusion with varying oxygenation. (B) Reconstructed photoacoustic (PA) images at 800 nm (upper), linear unmixed total haemoglobin (THb, middle), and linear unmixed blood oxygenation (sO^EST^, lower) maps. Image reconstruction artefacts can be observed near the blood inclusion and near the outer melanin layer (particularly clear in sO^EST^). (C) The photoacoustic signal in the outer cylinder of the phantom shows an increase with increasing melanin concentration. (D) Normalised photoacoustic spectra from the blood region show the effect of spectral colouring over a range of blood oxygenation levels. Scale bar = 5 mm.

Furthermore, examining the unmixed sO^EST^ as a function of the reference blood sO_2_ measurements reveals the effect of spectral colouring on PAI signal quantification (Figure 5 A, B), with higher melanin concentrations giving higher photoacoustic estimates of blood oxygenation when using linear spectral unmixing. Learned unmixing again reduces the disparity between low and high melanin concentrations, particularly at higher blood oxygenation levels (Figure 5 B, C). Linear unmixing estimates of sO^EST^ increase with melanin concentration (p *<* 0.001; gradient = 0.162 mL mg*^−^*^1^, 95 % CI [0.161 mL mg*^−^*^1^, 0.163 mL mg*^−^*^1^]), where as learned estimates do not (p = 0.50; gradient = 0.051 mL mg*^−^*^1^, 95 % CI [*−*0.603 mL mg*^−^*^1^, 0.706 mL mg*^−^*^1^]). Analysing simulations of digital twins of these phantoms in the tomographic PAI system showed consistent trends (Supplementary Figures S7, S8).

**Figure 5.**
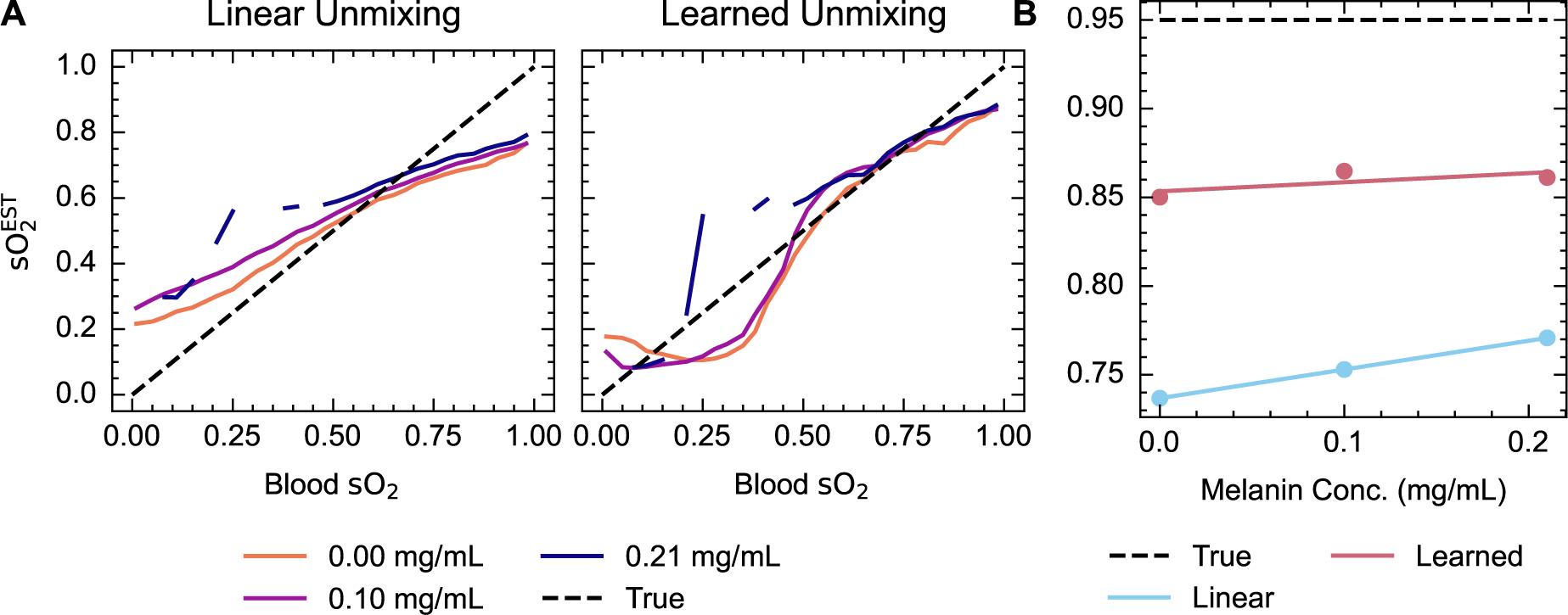
Comparison of linear unmixing and learned unmixing of blood oxygenation in skin-mimicking phantoms. Linear unmixing and learned unmixing blood oxygenation estimates (sO^EST^) are calculated and compared for varying melanin concentrations (A). Note that discontinuities in this plot arose from the rapid decrease in blood oxygenation from around 0.5 in the 0.21 mg mL*^−^*^1^. This is also shown as a function of melanin concentration at a high blood oxygenation, 95 % (B). The line of best fit from a linear model is shown. Linear unmixing sO^EST^ increases significantly (p *<* 0.001) with melanin concentration, while learned unmixing increases but not significantly (p = 0.50).

### 3.4 Mouse models underscore the severity of the impact of skin pigmentation on photoacoustics *in vivo*

Two closely related mouse models showing differential pigmentation were used to observe the effects of skin pigmentation on photoacoustic imaging *in vivo* (Supplementary Fig S3). Qualitative inspection of the image data clearly shows the increase in photoacoustic signal from the skin region in the presence of pigmentation and an associated decrease in photoacoustic signals measured from the body region in those pigmented mice (Figure 6 A, B). The spectra extracted from the ROIs show spectral colouring is clearly present in the pigmented mice. Spectra also demonstrate an increased presence of artefacts such as negative pixels (Figure 6 B). Line profiles across the mice (Figure 6 C-E) further emphasise these findings, highlighting the high level of optical absorption in the skin of the pigmented mice, artefactual increase in the total haemoglobin signals from the same region, and an increase in noise in the body of the mouse in the sO^EST^ image.

**Figure 6.**
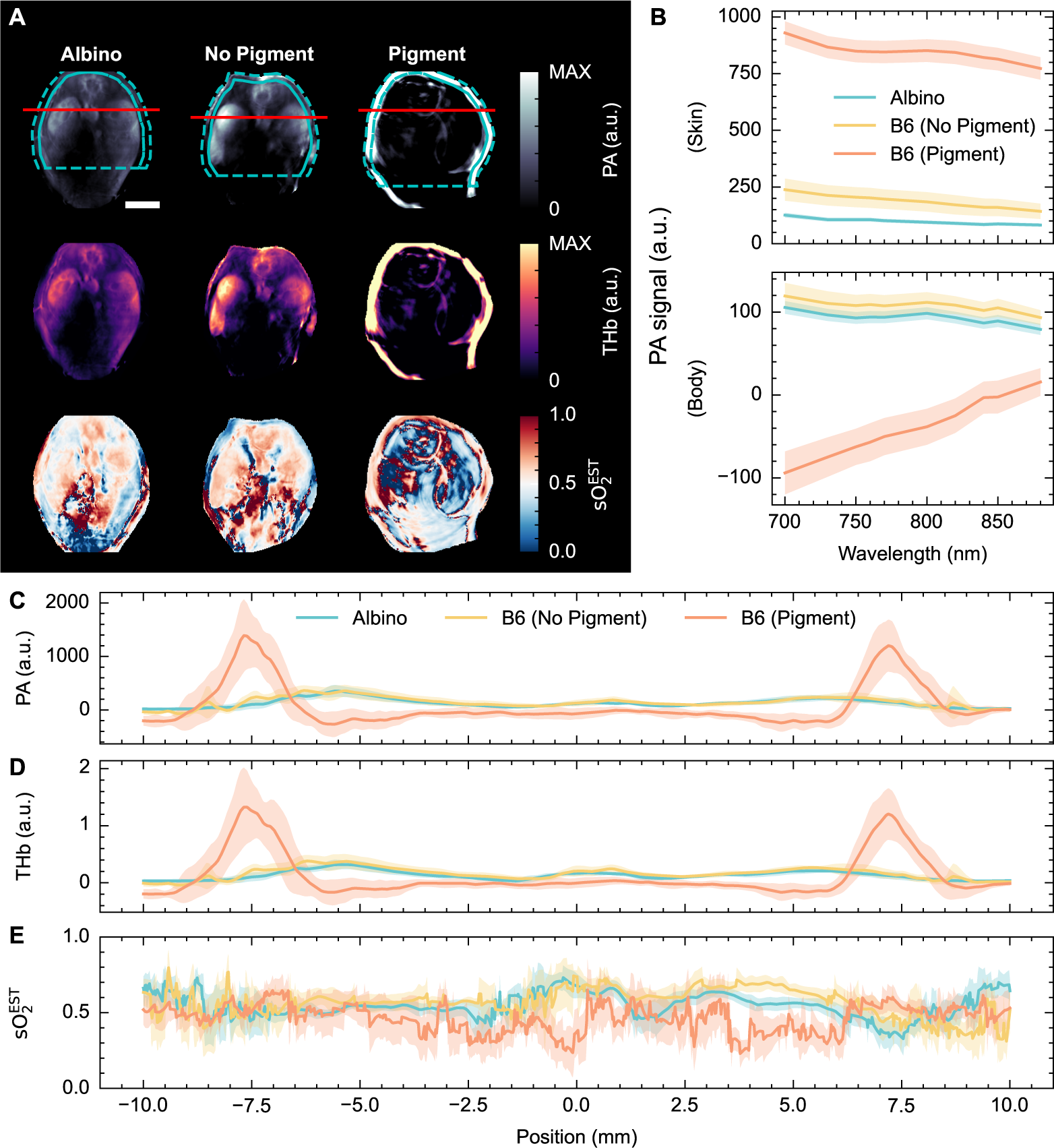
Mouse spleen and kidneys in albino and pigmented mice show substantial variations in photoacoustic imaging signal distribution. (A) Reconstructed photoacoustic images at 700 nm (upper), linear unmixed total haemoglobin (THb, middle), and linear unmixed blood oxygenation (sO^EST^, lower) maps. ROIs and line profiles used for quantitative analysis are shown on the single wavelength image. (B) Median photoacoustic spectra from the skin and body of albino mice (blue, n = 10), non-pigmented B6 mice (yellow, n = 8) and pigmented B6 mice (orange, n = 11). Line profiles through the kidneys, taken 6.5 mm from the surface of the mouse nearest the spine, of the photoacoustic signal at 700 nm (C), THb (D) and sO^EST^ (E). The shaded area shows the standard error. Scale bar = 5 mm.

Considering data from a single wavelength (700 nm, Supplementary Figure S9 A), the photoacoustic signal in the skin region of pigmented mice is significantly higher than in non-pigmented and albino mice (p = 0.010). Interestingly, there is also a significantly higher signal in the skin of non-pigmented B6 mice than in albino mice (p *<* 0.001). There was no significant difference between the photoacoustic signal, taken at 700 nm, in the body of the albino mice compared to the non-pigmented B6 mice (p = 0.95), however, there is a significantly lower signal in the body of the pigmented B6 mice (p *<* 0.001) (Supplementary Figure S9 B). The photoacoustic signal within the skin shows a negative correlation with the photoacoustic signal in the body of the mouse (p *<* 0.001) (Figure 7 A). Unmixed estimates of blood oxygen saturation are significantly higher (Figure 7 B) in pigmented mice compared to non-pigmented (p = 0.025) and albino mice (p = 0.019). There is a strong positive correlation between the skin photoacoustic signal and the sO^EST^ in the body of the mouse (Figure 7 C, p *<* 0.001), demonstrating that a higher melanin pigmentation in the skin leads to an overestimation of the underlying sO_2_.

**Figure 7.**
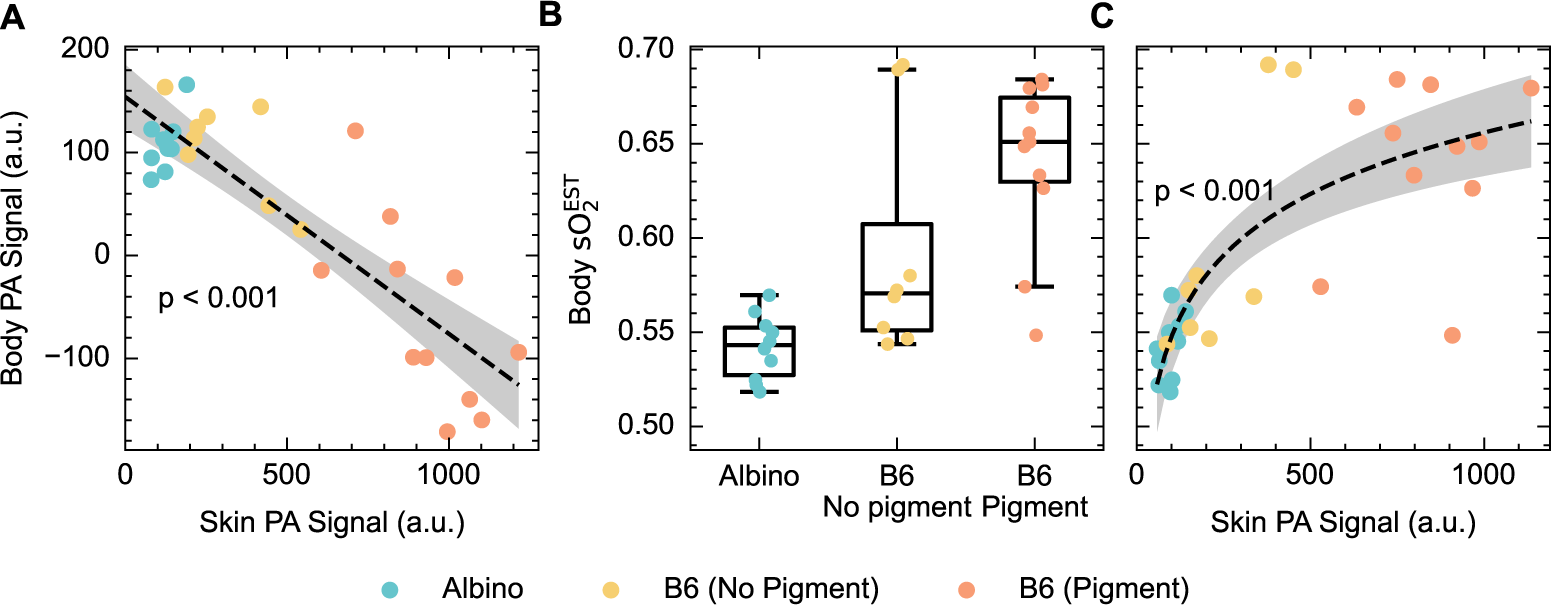
Increased skin pigmentation reduces the photoacoustic signal in the body and linear unmixing estimates of blood oxygenation are increased in pigmented mice. (A) Skin photoacoustic signal negatively correlates with body photoacoustic signal (p *<* 0.001). (B) The body sO^EST^ of B6 mice is significantly higher than that of albino mice (p = 0.025) and pigmentation is a significant factor in the linear model (p = 0.019). (C) Body sO^EST^ correlates with skin photoacoustic signal (p *<* 0.001).

Learned spectral unmixing was applied to the mouse data, with fair performance in albino and non-pigmented B6 mice. In pigmented mice, no clear improvement was observed over linear unmixing, with reconstruction artefacts dominating the oxygenation maps (Supplementary Figure S10).

## 4. DISCUSSION

Measurement bias in optical oximetry due to the differential light absorption of melanin in darker skin is widely acknowledged, yet the impact on PAI data has barely been tested. Tackling this challenge is key to ensuring equity in the use of photoacoustic imaging, and in biomedical optics more generally. To understand the underlying physics behind this phenomenon, we undertook a systematic study using a computational skin model, a simple tissue-mimicking phantom, and animals with differential skin pigmentation on the same genetic background. We then applied two different spectral unmixing approaches, linear unmixing and learned unmixing, revealing how spectral colouring caused by melanin can influence quantitative photoacoustic imaging.

Our findings confirm the presence of a measurement bias in PAI, with increased spectral colouring in the presence of melanin in all models. The spectral colouring trend is consistent with the optical properties of melanin; the absorption coefficient of melanin decreases with wavelength, so light propagation through tissue is more greatly attenuated at shorter wavelengths, a phenomenon that becomes more apparent as melanin concentration increases. Spectral colouring leads to a substantial overestimation of unmixed sO^EST^ with increasing Fitzpatrick skin type. The difference in sO^EST^ quantification between Fitzpatrick I and Fitzpatrick VI may be as large as 9 percentage points based on our simulations, which is similar to typical differences observed between cancer models (for example^54^). It is, therefore, unlikely that sO_2_ derived from linear spectral unmixing in its present form could be applied equitably when translated into the clinic for cancer detection or staging. The measurement of relative changes may be more robust to measurement biases, however, translation to the in vivo setting remains challenging, given the need for paired measurements and ground-truth data.

Fortunately, we observed that fundamental limitations in linear spectral unmixing, which have been characterised previously,^4, 49, 50^ may be partially mitigated using a learned spectral unmixing approach. Learned spectral unmixing demonstrated improved quantification relative to the ground truth in simulations and phantoms, and reduced the disparity between Fitzpatrick skin types, particularly in highly oxygenated blood. Learned unmixing, however, could not overcome the fundamental limitations of our backprojection-based reconstruction pipeline, which cannot accurately reconstruct the true initial-pressure distribution, particularly in regions adjacent to strongly absorbing structures. Strong optical absorbers lead to significant artefacts, such as streaking and negative pixels, as can be seen clearly in the centre of the images of heavily pigmented mice.^55^ Blood oxygen quantification was poor in negative pixel regions both with linear and learned unmixing.

Our study showed consistent agreement between Monte-Carlo simulations, phantoms, and in vivo studies, both in quantitative measures and image reconstructing artefacts. We expect our observations to transfer well into human data, however, there are further features of in vivo imaging that were not considered here and could further limit the use of photoacoustic imaging equitably in the clinic. In this study, we did not consider the effects of high acoustic impedance structures, such as bones, which could reflect the downward propagating photoacoustic wave from the epidermis, giving image reconstruction artefacts. Additionally, the effects of noise were not considered in our simulations, which would dominate in regions of low light fluence, such as deep in the tissue, particularly in the higher Fitzpatrick skin types. Additional noise could adversely affect the spectral unmixing results, particularly as the energy of different wavelengths will vary, and so affect the noise profiles. The presence of noise and acoustic reflection artefacts may explain the disagreement between this study and recently published *in vivo* data, which, contrary to this study, showed a decrease in linear spectral unmixing estimates of sO_2_ with increasing Fitzpatrick type.^29^ Future studies should explore ways to improve image reconstruction in the presence of strong optical absorbers and acoustic heterogeneity, for example with model-based reconstruction methods, and a substantial expansion of the available data from human subjects is required to thoroughly characterise this phenomenon across a range of PAI systems and methods.

## 5. CONCLUSION

PAI demonstrates a clear measurement bias with the presence of melanin in the skin. Spectral unmixing results were highly dependent on melanin concentration, which could introduce unintended biases into future human photoacoustic studies. Reproducible and equitable application of quantitative PAI will likely require a combination of physics-based, data-driven and empirical methods to account for varying light fluence and tissue properties fairly across the population.

## Disclosures

The authors have no conflicts of interest to declare.

## Code, Data, and Materials Availability

All data and code used in the preparation of this paper and not otherwise referenced in-line will be made available on the University of Cambridge Apollo repository (https://doi.org/10.17863/CAM.100220) or GitHub (https://github.com/BohndiekLab/melanin-phantom-simulation-paper) respectively.

## Acknowledgements

This work was funded by: Cancer Research UK (SB, TRE; C9545/A29580); the MedAccel program of the National Physical Laboratory financed by the Department for Business, Energy and Industrial Strategy’s Industrial Strategy Challenge Fund (LH); Cancer Research UK RadNet Cambridge [C17918/A28870] (EB); the Herchel Smith Studentship, Cambridge Trust (RT); and the Walter Benjamin Stipendium of the Deutsche Forschungsgemeinschaft (JG). We thank the Cancer Research UK Cambridge Institute Imaging Core and Biological Resources Unit for their support in conducting this research.

## Notes

### Competing Interest Statement

The authors have declared no competing interest.

