## Supplementary Materials for "The effects of skin tone on photoacoustic imaging and oximetry"

### Supplementary figures

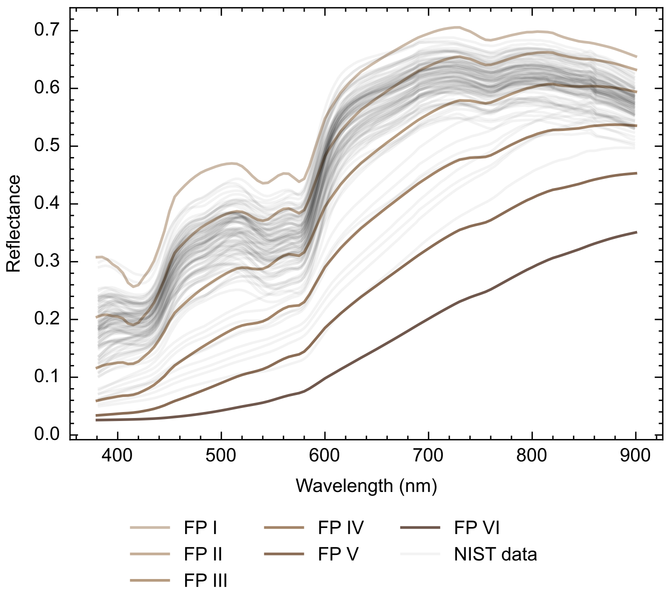


Figure S1. Simulated diffuse reflectance measurements (shades of brown, Fitzpatrick scale) are qualitatively consistent with integrating sphere measurements made of the forearm of volunteers in a publicly available dataset.[39]

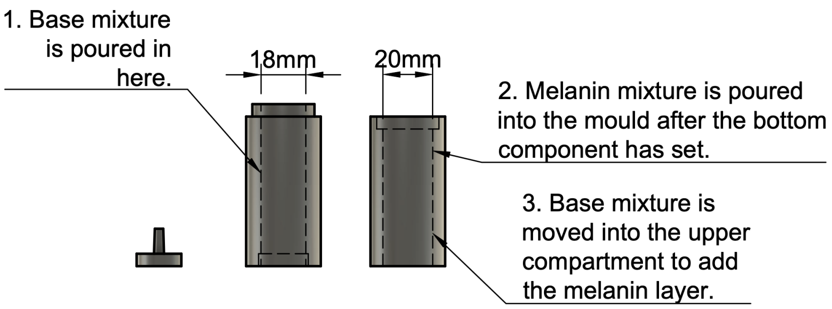


Figure S2. **Schematic diagram of the 3-D printed phantom mould.** A cross-section through the centre of the cylindrical mould. Left: the base of the mould, with Luer lock attachment for a needle and plastic tubing attachment, which is placed in the centre of the phantom. Right: a cross-section through the cylindrical mould, with a lower compartment in which the base mixture of the phantom is poured, and an upper compartment in which the melanin mixture is poured.

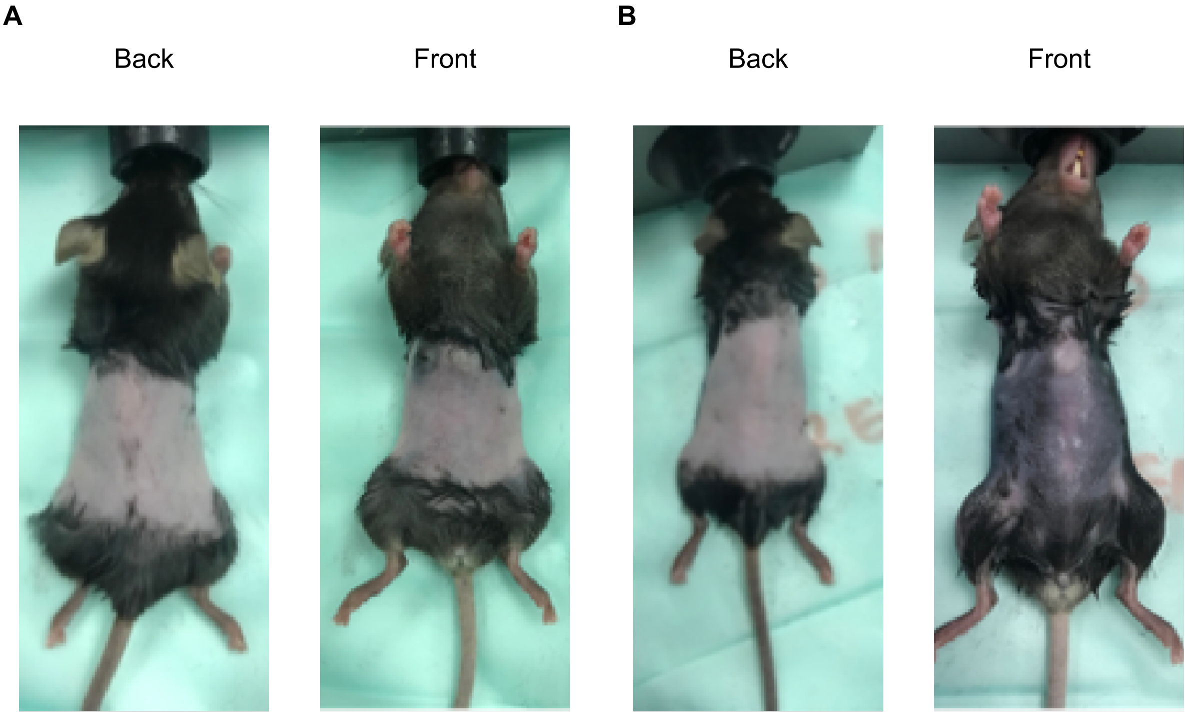


Figure S3. **Example image of B6 mice assigned to non-pigmented and pigmented group.** (A) An example of a non-pigmented mouse (prone and supine). (B) An example of a pigmented mouse (prone and supine.

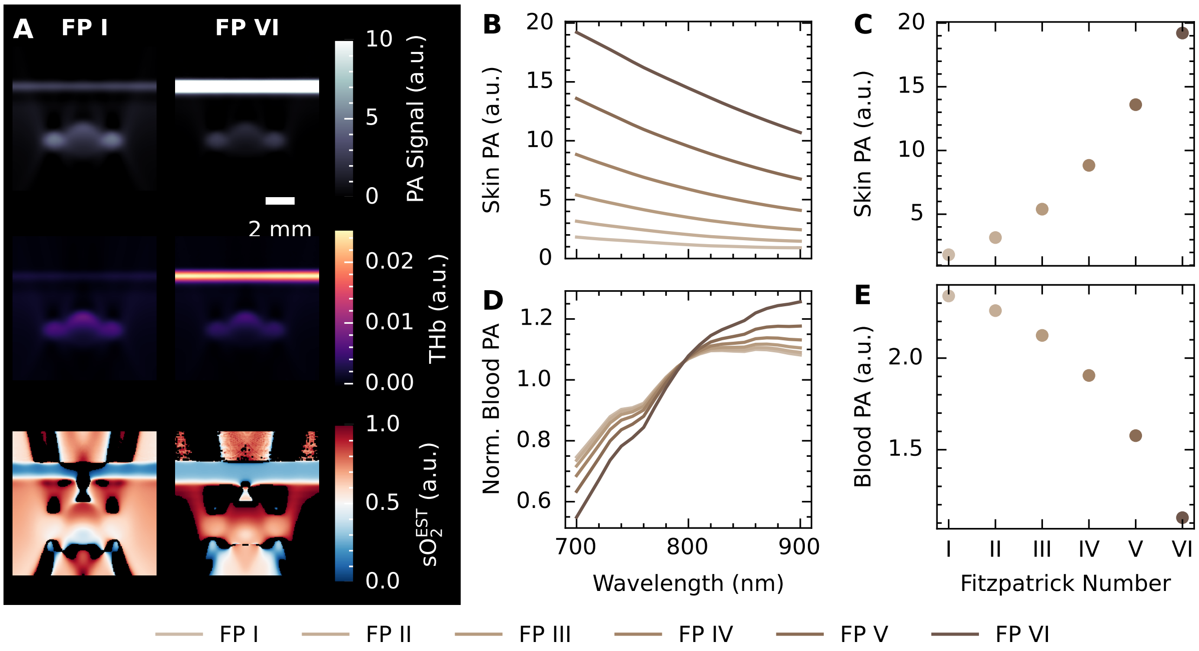


Figure S4. **Full optical and acoustic simulations show decreased signal and spectral colouring with increasing Fitzpatrick skin type, consistent with purely optical simulations.** (A) Photoacoustic reconstructed images, unmixed total haemoglobin (THb), and unmixed blood oxygenation ($sO_2^{EST}$) at each Fitzpatrick type for fully oxygenated blood in the forearm model. (B) Epidermis photoacoustic signal rises with increasing Fitzpatrick type at all wavelengths. (C) The photoacoustic signal at 700 nm increases with increasing Fitzpatrick type in the epidermis. (D) Increased spectral colouring is observed in the normalised blood photoacoustic spectra and (E) the photoacoustic signal at 700 nm decreases with increasing Fitzpatrick type in the blood vessel.

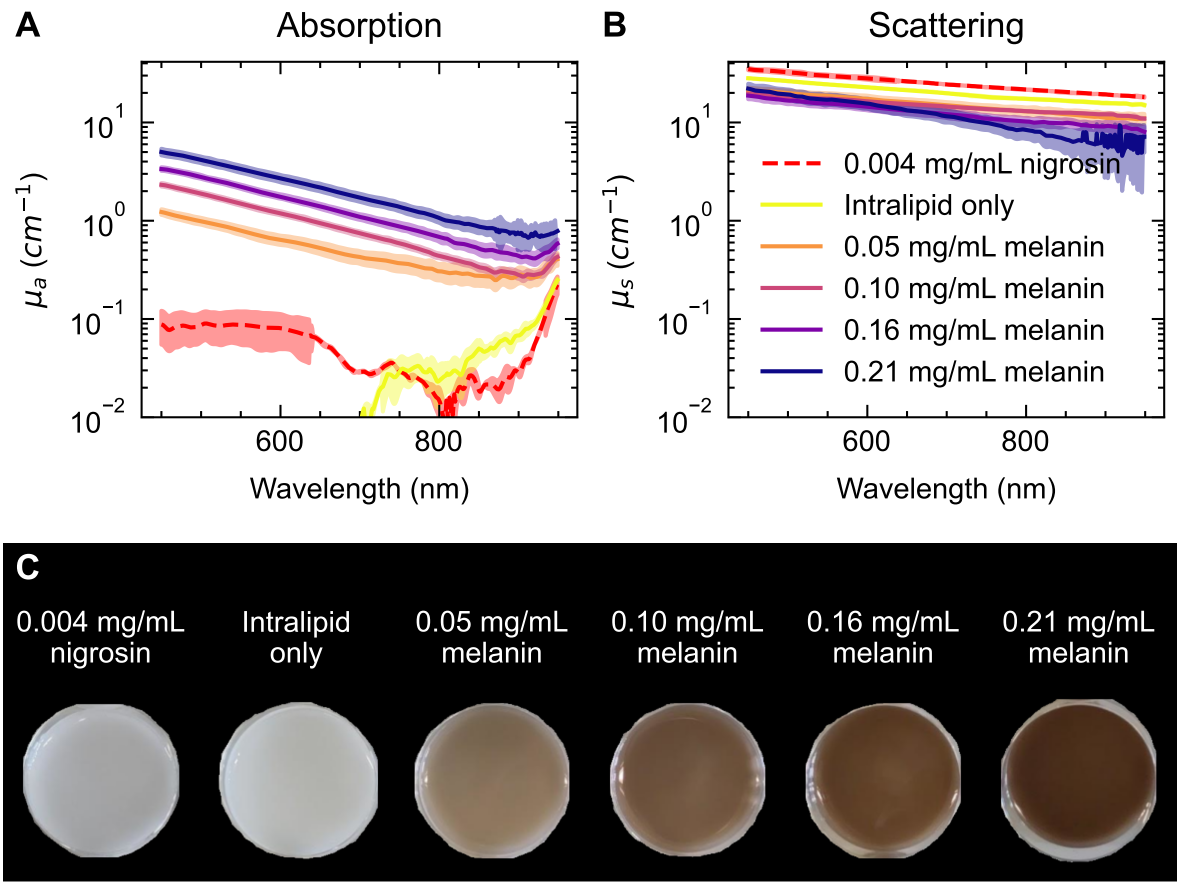


Figure S5. **Double integrating-sphere measurements confirm expected optical absorption and scattering properties of skin-mimicking phantoms.** Optical absorption (A) and scattering (B) coefficients as a function of wavelength were calculated using the inverse adding-doubling method. Shaded areas show 95 % confidence intervals. (C) Photographs of the skin-mimicking phantom material show qualitative agreement with the higher Fitzpatrick skin types.

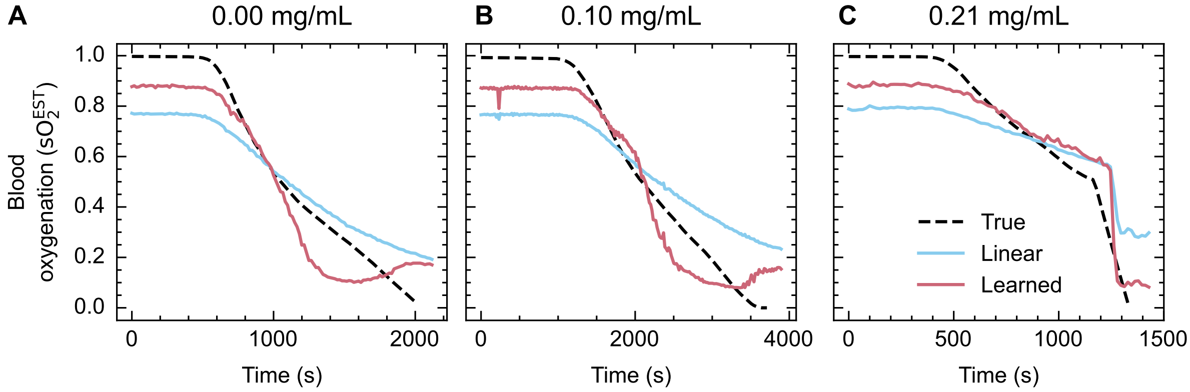


Figure S6. **Varying blood oxygenation in the phantom tubing over time.** The true blood oxygenation, as measured by a $pO_2$ probe and converted to $sO_2$ using the Severinghaus equation, is compared to the photoacoustic estimates of $sO_2^{EST}$ using linear unmixing and learned unmixing for each level of melanin concentration (A) $0.0\,mg\,mL^{-1}$ (b) $0.1\,mg\,mL^{-1}$ and (C) $0.21\,mg\,mL^{-1}$.

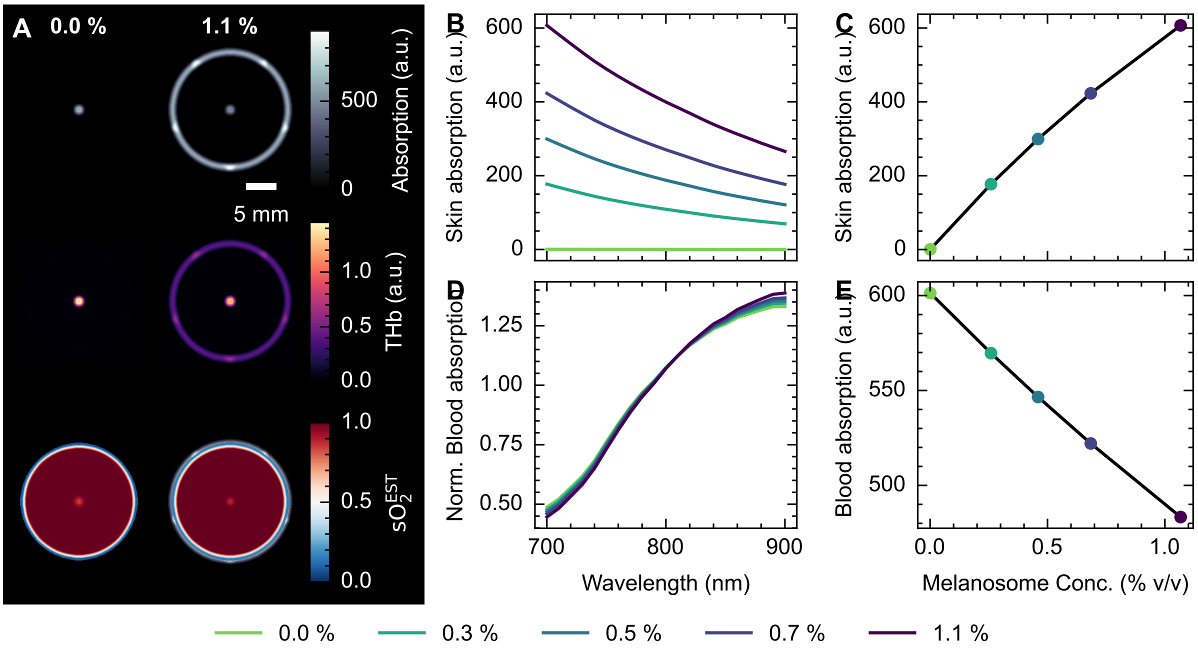


Figure S7. **Optical simulations of blood-flow phantoms reveal decreased blood signal with increasing melanin concentration and spectral colouring in the blood spectra.** (A) Initial pressure, and linear unmixing total haemoglobin and blood oxygenation, $sO_2^{EST}$ in representative simulations. (B) Initial pressure spectra of the melanin layer as a function of wavelength and (C) at 700 nm as a function of melanosome volume percentage. (D) Mean-normalised initial pressure spectra of the blood as a function of wavelength, and (E) non-normalised initial pressure spectra of the blood at 700 nm as a function of melanosome volume percentage.

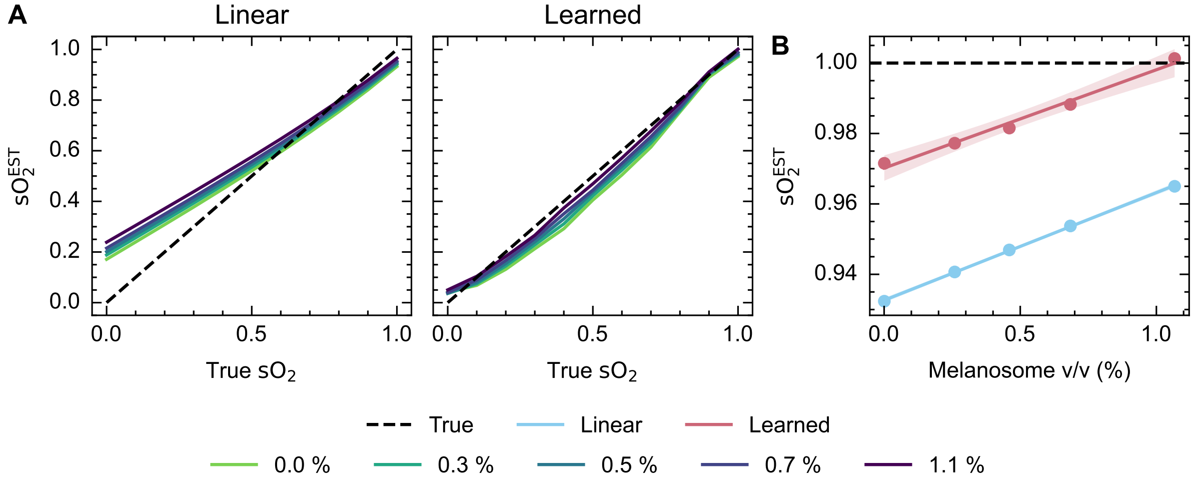


Figure S8. **Spectral unmixing results when applied to the optical simulations of a blood-flow phantom reveal biases due to spectral colouring.** Linear unmixing of the initial pressure distributions reveals a melanin concentration-dependent bias with both learned and linear unmixing (A). Colour scale corresponds to melanosome concentration. Photoacoustic blood oxygenation estimates increase with melanosome volume percentage in the outer layer (B).

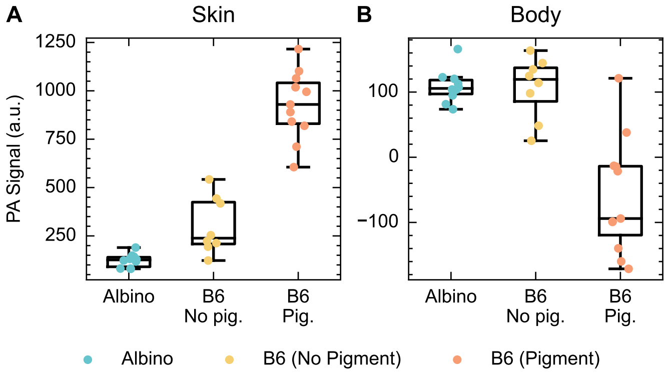


Figure S9. **Increased pigment in the mouse skin leads to increased photoacoustic signal in the skin and decreased photoacoustic signal in the body.** B6 mice have a significantly higher photoacoustic signal at 700 nm in the skin than albino mice (p=0.010) and pigmentation significantly increases this further ($p < 0.001$) (A). Non-pigmented B6 mice do not differ in their photoacoustic signal in the body ($p = 0.95$), however, pigmentation significantly reduces the photoacoustic signal in the body ($p < 0.001$) (B).

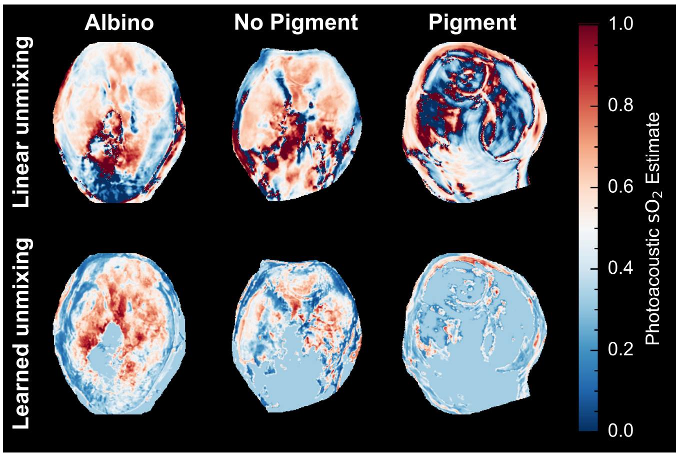


Figure S10. **Learned spectral decolouring performs poorly with *in vivo* mouse imaging data.** Linear unmixing (top) is compared to learned unmixing (bottom) in albino mice, non-pigmented B6 mice and pigmented B6 mice.

### Supplementary tables

Table S1. Tabulated values of the phantom parameters and approximate Fitzpatrick skin type, accounting for the thickness of the phantom melanin layer.

| Melanin concentration ($mg\,mL^{-1}$) | Absorption coefficient at 685 nm ($cm^{-1}$) | Estimated equivalent Fitzpatrick skin type |
|---|---|---|
| 0.00 | 0.00123 | Below I |
| 0.05 | 0.447 | II |
| 0.10 | 0.793 | III |
| 0.16 | 1.18 | III/IV |
| 0.21 | 1.84 | IV |
